# Predict future learning accuracy by the structural properties of the brain, an *in vivo* longitudinal MRI study in songbirds

**DOI:** 10.1101/477679

**Authors:** J. Hamaide, K. Lukacova, M. Verhoye, A. Van der Linden

## Abstract

Human speech and bird song are acoustically complex communication signals that are learned by imitation during a sensitive period early in life. Although the neural networks indispensable for song learning are well established, it remains unclear which neural circuitries differentiate good from bad song copiers. By combining *in vivo* structural Magnetic Resonance Imaging with song analyses in juvenile male zebra finches during song learning and beyond, we discovered that song imitation accuracy correlates with the structural architecture of four distinct brain areas, none of which pertain to the song control system. Furthermore, the structural properties of a secondary auditory area in the left hemisphere, are capable to predict future song copying accuracy, already at the earliest stages of learning, before initiating vocal practicing. These findings appoint novel brain regions important for song learning outcome and inform that ultimate performance in part depends on factors experienced before vocal practicing.

## Introduction

Human speech and bird song are highly complex and rapid motor behaviors that are learned by imitation and serve to produce complex communication signals vital for social interactions(1). Both are acquired during a sensitive period early in life which consists of a sensory learning phase where a perceptual target of speech sounds or a song model are memorized, followed by a sensorimotor learning phase where human infants or juvenile birds engage in intense vocal practicing and will try to match their own vocalizations to the previously established perceptual targets based on multi-sensory feedback(2).

In zebra finches, being skilled vocal learners, the success of the song learning process is quantified by computing the acoustic similarity between the pupil and the tutor song(3). That is, how accurate the pupil is capable of imitating the tutor song. This way, the quality of the imitation can only be assessed when birds have engaged in vocal practicing and, as such, is always a product of both sensory and sensorimotor learning. Importantly however, like human speech, bird song is a complex socially-learned communication signal and how successful pupils become at copying the tutor song also depends on the way the tutor interacts with the juvenile learning birds(4). Likewise, studies have shown that live tutoring, i.e. where the pupil bird is housed together with an adult male zebra finch, results in higher song similarity scores compared to tape tutored birds(5). As such, various aspects including social factors determine the final song learning accuracy of a juvenile bird. Even though the pathways necessary for both sensory and sensorimotor learning to succeed are relatively well characterized (Supplementary Fig. SI-1), it is still unclear which brain areas are engaged in determining the successfulness of the song learning process or, in other words, the song imitation accuracy. To address this question, one needs appropriate research tools that enable to repeatedly and over extended time frames capture the structural or functional properties of the entire zebra finch brain, when birds progressively reach excellent vocal imitation accuracy levels. Only recently, such methods, i.e. *in vivo* Magnetic Resonance Imaging (MRI), have become available to investigate the juvenile zebra finch brain(6).

In humans and small animal species, MRI emerged as a prominent tool to quantify adaptations to brain function and structure that arise along training or learning paradigms(7-9), or even to uncover novel and unexpected brain areas involved in a specific function(10) or task. Inspired by these imaging studies, we set up a longitudinal *in vivo* MRI study to expose brain areas that display structural changes when zebra finch males progressively advance from sensorimotor song learning towards post critical period song refinement. Exploiting the advantages of *in vivo* MRI, we performed brain-wide voxel-wise statistical analyses to quantitatively explore gross anatomy (3D, volume) or the intrinsic tissue properties (Diffusion Tensor Imaging (DTI)) of the entire zebra finch brain along the different stages of the song learning process. We discovered that song imitation accuracy correlates with the structural architecture of four distinct brain areas, none of which pertain to the song control system. Interestingly, the data reveal that the structural properties of secondary auditory region NCM, of the left hemisphere, are capable to predict future song learning success.

## Results

### Song performance improves even after crystallization

Sensorimotor song learning includes a gradual matching of the spectro-temporal characteristics of own vocalizations to acoustically match those of the target song(11). Reaching the end of the critical period for vocal learning, the song crystallizes indicating that in normal circumstances the learned song will remain unchanged. To quantitatively assess fine-scale changes in song performance and estimate song learning accuracy, we evaluated the progression of song learning from advanced sensorimotor (65 days post hatching (dph)), over the crystallization phase (90-120 dph), to fully crystallized song (200 dph) using three measures. First, we extracted the acoustical properties of individual song syllables at each age to evaluate how the spectral and temporal structure of the syllables evolve from (advanced) plastic to fully mature stereotyped song. These analyses informed that the syllables gain tonality (lower acoustic noisiness), become more structured (less noisy) and less acoustically diverse along sensorimotor song maturation (full description of the song analyses can be found in Supplementary Data 1). Furthermore, while the spectral content of syllables mainly forms during the sensorimotor phase, the temporal properties of the songs continue to change beyond the crystallization phase. This corroborates previous studies in zebra finches(12). Second, we quantified how the syllable sequence stabilizes towards song crystallization (sequence stereotypy), and, third, how accurate the bird is able to produce an acoustic copy of the tutor song (% similarity). We tested for age-related changes in performance or whether the tutor takes part in the juveniles’ learning success (summary of results: Supplementary Data 1).

Song crystallization results in a highly stereotyped, consistent order of syllables within a motif. This is reflected in sequence stereotypy which increased gradually from 65 to 200 dph (*p*=0.0052 *F*_*(3,38.4)*_ =4.7904, Figure 1B). To quantify how successful the juvenile birds learned, i.e. copied, the tutor song, we computed the spectral similarity to the tutor song. Song similarity to tutor song increased gradually from 65 to 200 dph (*p*=0.0251 *F*_*(3,37.0)*_=3.4890; Figure 1A) reaching similar levels as described by others(3, 13). Interestingly, both song sequence stereotypy and song similarity to tutor song progressively increase from 65 to 200 dph suggesting that even ‘crystallized’ song is still refined to achieve a better acoustic match with the previously memorized tutor song. Importantly, the overall variability in song similarity between birds is quite large indicating that not all birds copy the tutor song equally well. Prior studies have shown that successful song learning not only depends on the ability of the juveniles to hear the song model. Social interactions between the juvenile and tutoring bird are crucial mediators in determining the overall quality of the song copy(4). Likewise, we observe that song similarity depends on tutor identity, that is, juveniles raised by a specific tutor will consistently sing a good or less good tutor song copy (*p*=0.0159 *F*_*(7,6.1)*_=6.7597). This main effect of tutoring bird on song similarity outcome did not seem to depend on the length of the songs, as based on similarity scores obtained at 200 dph no differences in the number of syllables for birds having a ‘high’ (>68%) or ‘low’ (<68%) similarity score could be observed (Supplementary Data 1 Table SI-I-B).

**Figure 1:**
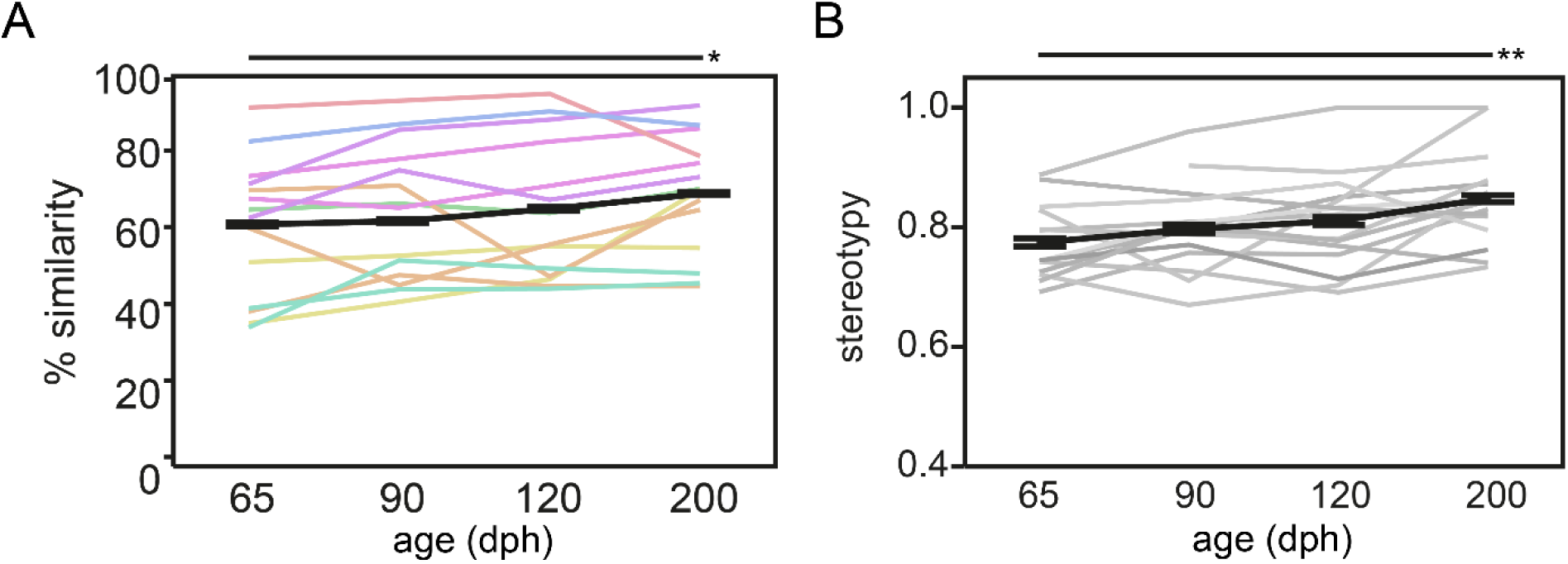
Song similarity improves beyond crystallization. Each thin colored or grey line refers to the average performance of an individual bird over the different ages. The bold black line presents the average group performance (mean ± s.e.m.; n=14; 20 data points per time point per bird). Graph **A** and **B** refer to respectively song similarity to tutor song and song stereotypy in function of age. Both increase from 65 to 200 dph (mixed-effect model main effect of age: song similarity: *p*=0.0251 *F*_*(3,37.0)*_=3.4890; sequence stereotypy: *p*=0.0052 *F*_*(3,38.4)*_=4.7904). The color-code of the lines in A encodes tutor identity, i.e. birds raised by the same tutor share the same color. The color-code illustrates that song similarity is dependent on tutor identity (mixed-effect model main effect for tutor: *p*=0.0159 *F*_*(7,6.1)*_=6.7597). Asterisks indicate significant differences over time identified by a mixed model analysis with *post hoc* Tukey’s HSD test. *: p<0.05; ** p<0.01.

### Imitation accuracy traces back to the CM, VP, tFA and NCM

Even though song performance improves from the sensorimotor to the crystallization phase (and even beyond), not all birds learn the tutor song equally well. Therefore, we set out to explore whether better song performance, i.e. more accurate song copying, coexists with a specific structural signature in the brain. Inspired by ample *in vivo* imaging studies describing training- or learning-induced brain-behavior relationships(8), we performed brain-wide voxel-based statistical analyses to highlight potential brain areas that present a correlation between song learning accuracy (% similarity between pupil and tutor song) and local volume or intrinsic tissue properties derived from the DTI metrics (Figures 2-3-4, Supplementary Data 2 Table SI-2-A). These analyses uncovered four clusters. More specifically, the clusters co-localize with two secondary auditory areas, i.e. the caudomedial nidopallium (NCM) and caudal mesopallium (CM), with a white matter tract that connects the basorostral nucleus to the arcopallium (frontoarcopallial tract (tFA)(14)), and an area at the base of the telencephalon termed the ventral pallidum (VP). The VP contains many fibers of passage among those that project from the midbrain dopaminergic nuclei to Area X of the anterior forebrain pathway and Area X-DLM projections(15). Surprisingly, none of these areas pertains to the traditionally studied song control system.

**Figure 2:**
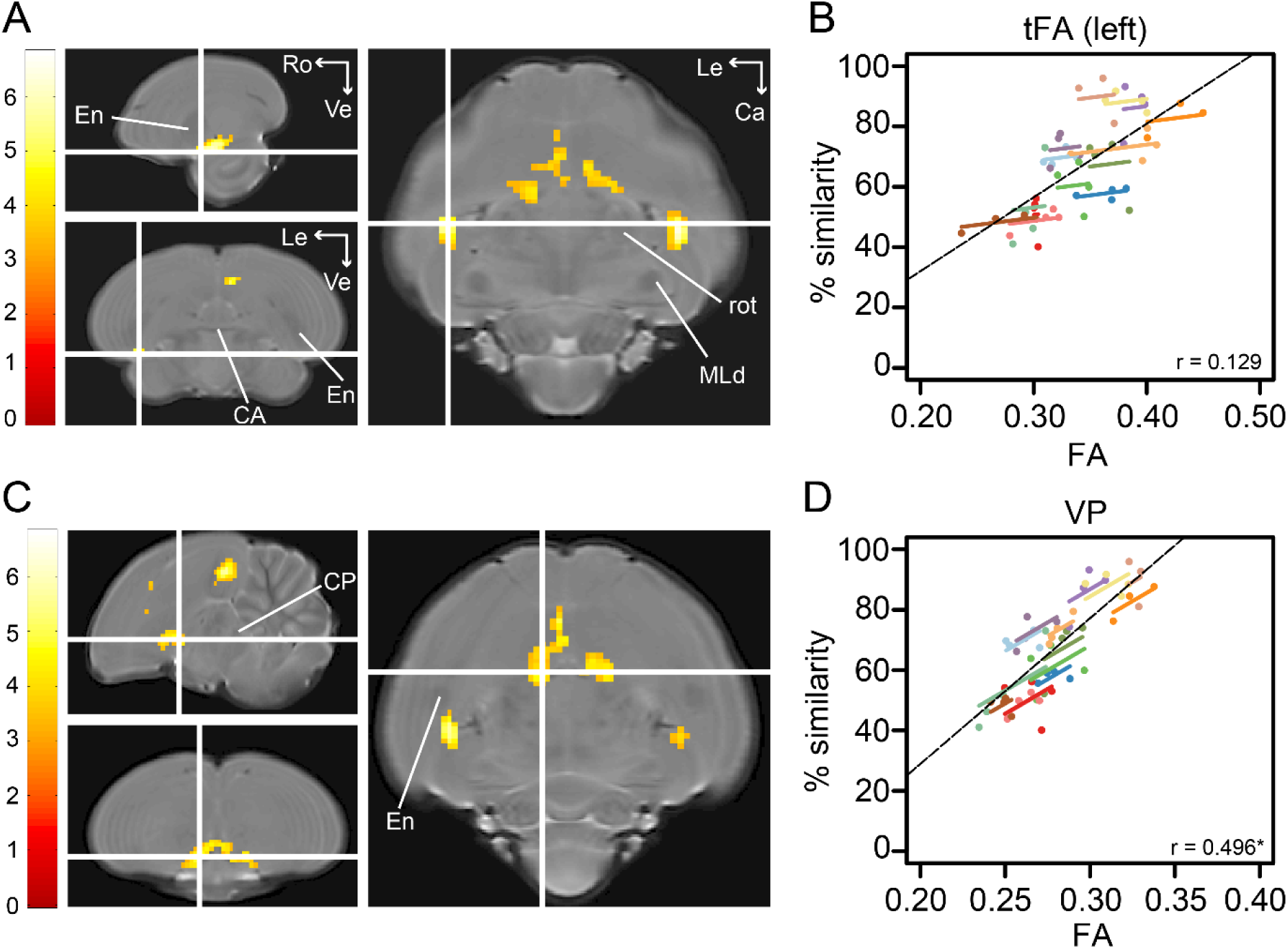
Song imitation accuracy correlates positively with Fractional Anisotropy in the tFA (A-B) and the VP (C-D). The statistical maps (A and C) present the outcome of the voxel-based multiple regression testing for a correlation between song similarity and FA (n=14). The crosshairs point to the tFA in the left hemisphere (A) or the VP (C). Results are overlaid on the population-based MRI template and scaled according to the color-code on the left (T values). Only voxels that reached _*puncorrected*_<0.001 and take part of a cluster of at least 40 contiguous voxels are displayed. Graphs B and D visualize the nature of the correlation between song similarity and FA where individual data points are color-coded according to bird-identity (i.e. one color = one bird). The average within-bird correlation is presented by the colored lines, while the black dashed line indicates the overall association between song similarity and FA, disregarding bird-identity or age. ‘r’ is the repeated-measures correlation (rmcorr) coefficient. The % indicates a significant rmcorr correlation between FA and * similarity in the VP (*p*=0.001). Abbreviations: CA: anterior commissure; CP: posterior commissure; En: entopallium; MLd: dorsal part of the lateral mesencephalic nucleus; rot: nucleus rotundus; tFA: fronto-arcopallial tract; VP: ventral pallidum.

The voxel-based analyses identified a significant positive correlation between % similarity and Fractional Anisotropy (FA) in the left tFA (peak: *p*_*FWE*_<0.001 *T*=6.81; Figure 2 A), and in left NCM (rostral NCM; peak: *p*_*FWE*_=0.019 *T*=5.69; Figure 3 A). FA quantifies the directional dependence of water diffusion(16). As a result, alterations to FA can be caused by a wide variety of microstructural tissue re-organizations including altered axonal integrity, myelination, axon diameter and density, change in cellular morphology, etc.(8, 16). Furthermore, we uncovered an additional cluster midsagittal near the striatum and mesopallium, extending laterally and caudo-ventrally adjacent to the septomesencephalic tract (TSM; sub-peak next to the TSM in the left hemisphere: *p*_*FWE*_=0.002 *T*=6.38; Figure 2 C). Based on this spatial pattern and in accordance with the Karten-Mitra zebra finch brain atlas(17), we identify this area as the VP. Interestingly, when inspecting the statistical maps at an exploratory threshold (*p*_*uncorrected*_<0.001 k_E_≥40voxels), clusters could be observed at the right tFA (peak: *p*_*FWE*_=0.001 *T*=6.42) and the right NCM (peak: *p*_*FWE*_=0.032 *T*=5.55; Figure 3 C) as well. The cluster covering the left NCM extends rostro-laterally towards the CM (sub-peak of NCM cluster: *p*_*FWE*_=0.194 *T*=5.01 Figure 3 inset B_1_).

**Figure 3:**
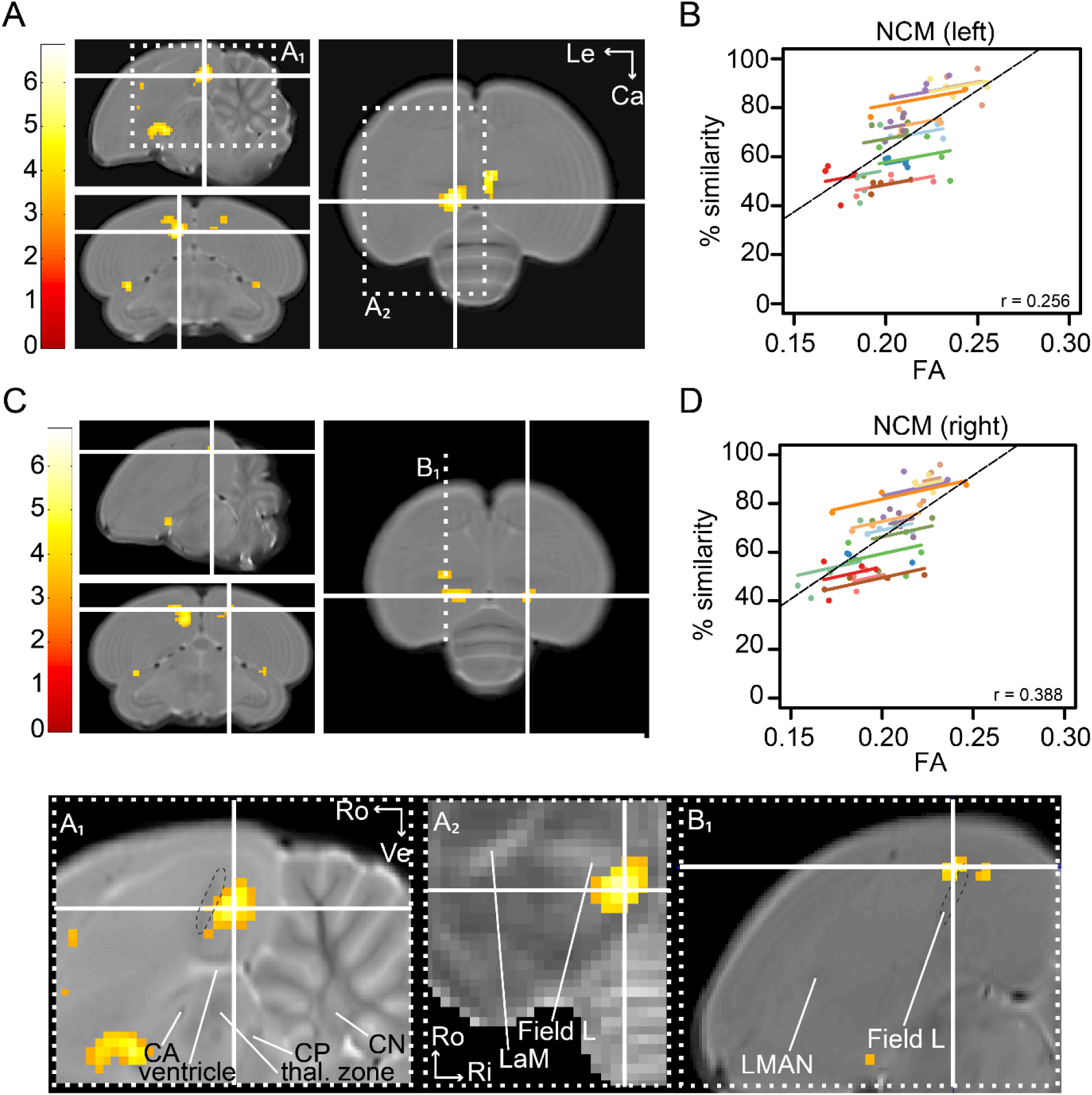
Song imitation accuracy correlates positively with Fractional Anisotropy in NCM. The statistical maps present the outcome of the voxel-based multiple regression (n=14) testing for a correlation between song similarity and FA. The crosshairs converge in the NCM of the left (A) and right (C) hemisphere. Results are overlaid on the population-based template or population-averaged FA maps of datasets obtained at 200 dph (A_2_), and are scaled according to the color-code on the left (T values). Only voxels that reached *p*_*uncorrected*_<0.001 and take part of a cluster of at least 40 contiguous voxels are displayed. Insets A_1-2_ visualize the spatial extent of the cluster in left NCM with reference to Field L. L2a is visible by a hypo-intense line-shaped area on A_1_ (black dashed circle). The peak of the cluster (crosshairs) is situated in (rostral) NCM. Inset A_2_ further corroborates that the NCM-cluster does not overlap with the highly myelinated, fiber-rich (hyper-intense FA) structure that can be identified as Field L. Inset B_1_ illustrates the cluster in left NCM that extends to the left CM (rostral to Field L (black dashed circle). Graphs B and D visualize the nature of the correlation between song similarity and FA, in NMC of the left and right hemisphere respectively, where individual data points are color-coded according to bird-identity (i.e. one color = one bird). The average within-bird correlation is presented by the colored lines, while the black dashed line indicates the overall association between song similarity and FA, disregarding bird-identity or age. ‘r’ is the repeated-measures correlation (rmcorr) coefficient. Abbreviations: CA: anterior commissure; CP: posterior commissure; LaM: *lamina mesopallialis*; LMAN: lateral magnocellular nucleus of the anterior nidopallium; thal. zone: thalamic zone.

The voxel-based multiple regression uncovered a significant negative correlation between % similarity and local tissue volume in the VP (peak: *p*_*FWE*_<0.001 *T*=8.06; Figure 4 A). In addition, another bilateral cluster displaying a negative correlation between local volume and % similarity was observed rostral to Field L in the medial and lateral CM (CMM and CLM, respectively), potentially including nucleus avalanche (Av) (left: peak: *p*_*FWE*_=0.001 *T*=7.10; right: peak: *p*_*FWE*_<0.001 *T*=7.42; Figure 4 C).

**Figure 4:**
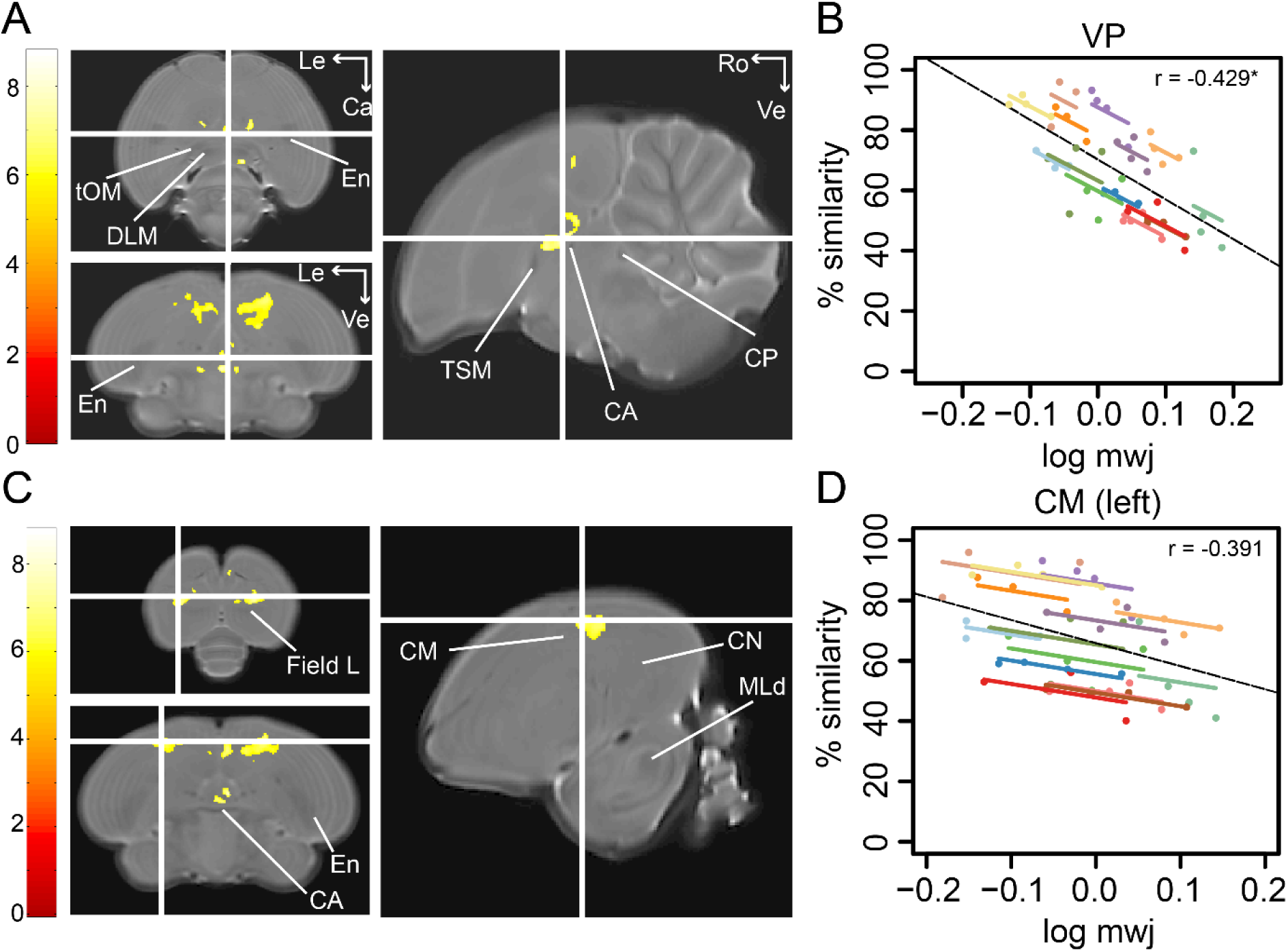
Song imitation accuracy correlates negatively with the local volume of the VP (A-B) and the CM (C-D). The statistical parametric maps present the outcome of the voxel-based multiple regression testing for a correlation between song similarity and local tissue volume (n=14) are visualized at *p*_*FWE*_<0.05 and k_E_≥80 voxels, and overlaid on the population-based template. The crosshairs point to the VP (A) or the CM in the left hemisphere (C). T-values are color-coded according to the scale immediately next to the SPMs. Graphs B and D inform on the nature of the association between song similarity (%) and log-transformed modulated jacobian determinant (log mwj; a metric reflecting local tissue volume). More specifically, the individual data points of the graphs are color-coded according to bird-identity (i.e. one color = one bird). The average within-bird correlation is presented by the colored lines, while the dashed black line indicates the overall association between song similarity and log mwj, disregarding bird-identity or age. ‘r’ is the repeated-measures correlation (rmcorr) coefficient. The * indicates a significant rmcorr correlation between logmwj and * similarity in the VP (*p*=0.0057). Abbreviations: CA: anterior commissure; CM: caudal mesopallium; CN: caudal nidopallium; CP: posterior commissure; DLM: medial part of the dorsolateral nucleus of the anterior thalamus; En: entopallium; MLd: dorsal part of the lateral mesencephalic nucleus; tOM: occipitomesencephalic tract; TSM: septo-mesencephalic tract; VP: ventral pallidum.

### Learning-related relationships versus between-bird variance

The voxel-based multiple regression detects an overall association between song performance (% similarity) and the structural properties of the brain without taking the repeated measures into account. As a result, these analyses cannot inform whether the brain-behavior associations are mainly driven by between-subject variation in performance and structure, or whether individual improvements in song imitation as a direct consequence of the sensorimotor learning or post-crystallization song refinement process relate to specific structural properties of the clusters identified by voxel-based analyses. To make this distinction, we first extracted for each bird and each time point separately the mean log-transformed modulated jacobian determinant or mean FA from the voxel-based clusters. Next, we performed (1) Spearman’s correlation analysis (ρ) to characterize potential correlations between the structural properties of the cluster-based ROIs and song similarity at 65 or 200 dph, and (2) a repeated-measures correlation analysis (rmcorr(18)) which informs on the common within-bird association between brain structure and song similarity across the group of birds including data of all four time points. The rmcorr analysis reveals whether on average across birds individual improvements in song learning accuracy (% similarity) correlate with increased FA or decreased volume. In other words, this analysis informs on potential learning-related changes in local brain structure.

The results of the correlation analyses are summarized in Supplementary Data 3 and visualized in Figure 2 B and D, Figure 3 B and D, and Figure 4 B and D. The voxel-based association between song similarity and FA in the NCM and in the tFA appears to be driven by between-subject variance (65 dph: NCM: left: *p*=0.0081; right: *p*=0.0138; tFA: left: *p*= 0.0009; right: *p*=0.0045; 200 dph: NCM: left: *p*=0.0070; right: *p*=0.0336; tFA: left: *p*= 0.0012; right: *p*=0.0041). Surprisingly, the small cluster in the right NCM displays a significant repeated-measures correlation indicating that when individual birds improve their performance, FA increases accordingly. In contrast, the CM presents no between-subject correlations at any age. However, individual improvements in song learning result in a lower local volume of the CM (left: *p*=0.0126; right: *p*=0.0075). The VP presents significant repeated-measures correlations between song similarity and local volume or FA (log mwj: *p*=0.0057; FA: *p*=0.0010). This informs that when individual birds learn to produce a more accurate copy of the tutor song, the VP scales down and obtains a more ordered structure. Furthermore, also between birds, local volume and FA correlate significantly with song similarity during the sensorimotor phase (log mwj: *p*=0.0006; FA: *p*=0.0004), however, only the correlation between song similarity and FA is maintained until after song crystallization (*p*=0.0008).

Together, these findings suggest that birds that sing a better copy of the tutor song have higher FA values in NCM, the VP and the tFA, both during the sensorimotor phase and after crystallization. Furthermore, learning-related individual advances in producing a more accurate acoustic copy of the tutor song correlate with local tissue structure in the caudal mesopallium and VP. Overall, higher song similarity is related to a smaller volume of the VP and CM or a higher diffusion anisotropy (more ordered structure) in the VP, tFA and NCM. Higher anisotropy might refer to a more accurate alignment of fibers in the VP and tFA, while in grey matter-like structures such as NCM, higher FA values might allude to changes in cell morphology (spines and dendrite branching) or density, etc.(8).

### NCM predicts future good or bad learning outcome

The Spearman correlation analyses uncovered that FA values in the VP, NCM and tFA present a clear between-subject correlation with song learning accuracy. This suggests that in the sensorimotor phase good or bad learners are characterized by a distinct structural MRI parameter readout in these regions. Next, based on the scans of the same birds acquired during the sensory (20, 30 dph) and early sensorimotor (40 dph) phases, we evaluated whether similar signs of future good or bad song copying outcome would already be visible in the structural properties of these regions in the early sensorimotor phase, or even before sensorimotor practicing, during the sensory learning phase, when birds memorize the tutor song but are not yet fully engaged in trial-and-error vocal practicing(1).

We divided the group of male birds into ‘good’ and ‘bad’ learners based on the overall song performance obtained at 65-200 dph. More specifically, good learners (n=7) always sung acoustically accurate copies (>65-68% song similarity to tutor song), while bad learners (n=5) never produced a copy better than 65-68% similarity to tutor song. Birds that clearly traversed the 65-68% interval throughout the study (n=2) were not considered in this analysis (Figure 5 A). Next, we tested for an interaction between age (20-30-40 dph) and future learning accuracy (good, bad) in the cluster-based ROIs (Supplementary Data 4, Figure SI-3). None of cluster-based ROIs survived FDR correction for multiple comparisons when testing for an interaction between age and future learning accuracy. In contrast, FA in the left NCM displayed a significant main effect of imitation accuracy in males aged 20-40 dph (*p*=0.0003 *F*_*(1,10.2)*_=28.3681; Figure 5 B-C). Good learners consistently demonstrate higher FA values in left NCM compared to bad learners. Intriguingly, this difference in FA is already present at 20 dph, that is, before the start of the sensory learning phase(19) when the juvenile finches have been exposed to their tutor for several days.

**Figure 5:**
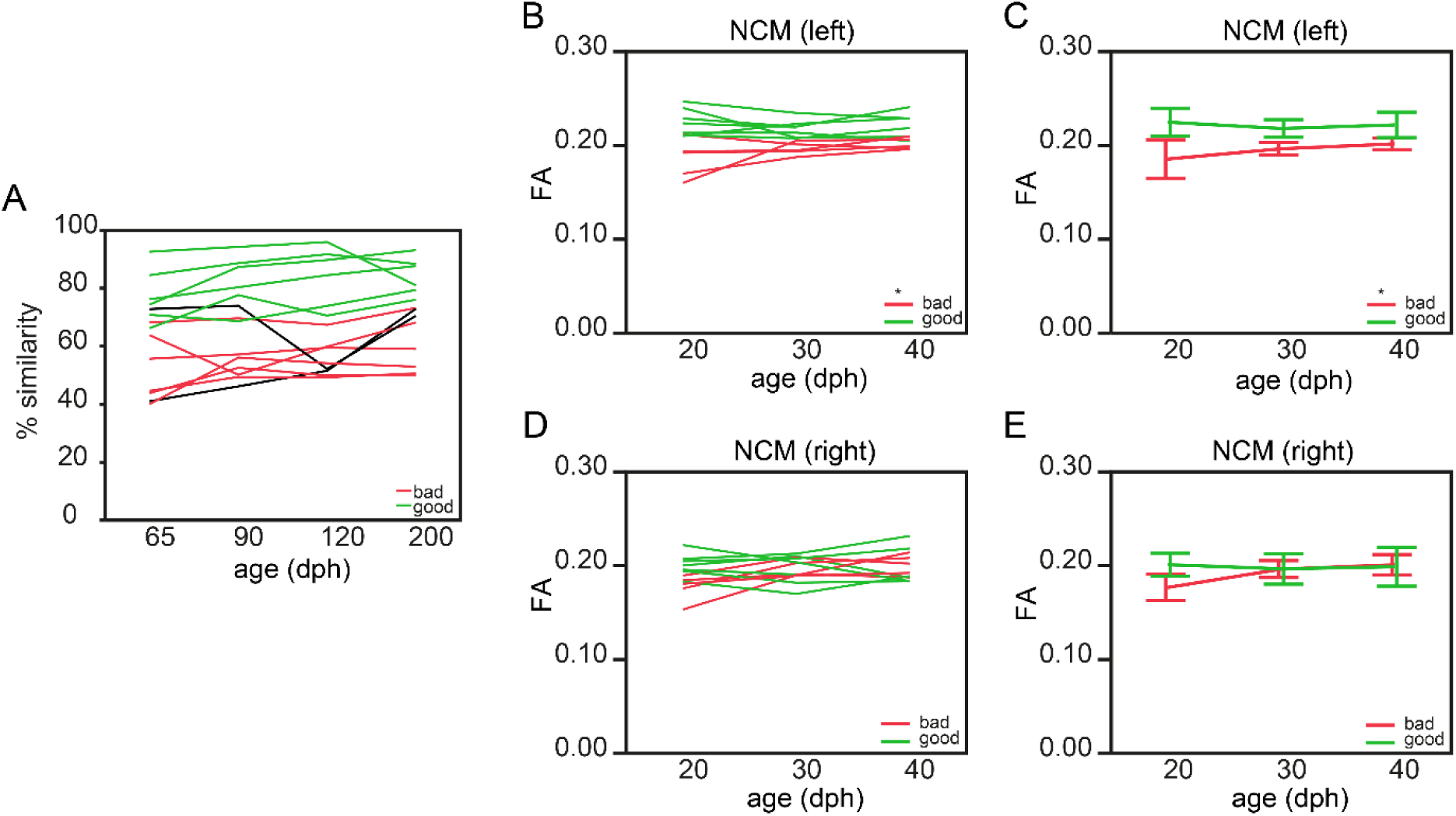
Fractional Anisotropy in left NCM predicts future song learning accuracy. Graph A presents the learning curve of the good (green) and bad (red) learning birds from 65 to 200 dph. The two birds that clearly traversed the 65-68% song similarity score could not be assigned to either group and are presented by a thin black line. (Details on the distinction between good and bad learners can be found in the Results section.) Graphs B-E present the difference of FA in NCM between good (green) and bad (red) vocal learners during the sensory (20-30 dph) and early sensorimotor (40 dph) phase in both hemispheres separately. In B and D one line represents one bird, while C and E provide the group mean ± standard deviation. The * indicates a significant main effect between good and bad learners in left NCM (mixed model: *p*=0.0003 *F*_*(1,10.2)*_=28.3681).

## Discussion

The major aim of this study was to expose brain regions that are actively shaped along the process of song learning in male zebra finches, when –as practice makes perfect– pupils progressively become more proficient at imitating the tutor song. We employed recently implemented brain-wide *in vivo* structural imaging tools(6) and used song similarity to tutor song as a proxy for vocal learning accuracy, starting from the advanced sensorimotor phase up to post critical period song refinement. This resulted in two interesting findings. First, we discovered that the structural properties of the secondary auditory cortices, i.e. left NCM, the CM, and unexpectedly, the VP and the tFA correlate with imitation accuracy. Between- and within- subject correlation analyses revealed that the structural properties of left NCM and the tFA are mainly caused by between-subject variation in performance and structure, while the structural architecture of the CM and VP appears to change along the learning process, in individual birds. Second, we demonstrate that the structural properties of left NCM differ between future good or bad song learning outcome already during the sensory phase, when birds establish a memory of the tutor song but have not yet initiated sensorimotor vocal practicing. As such, the structural properties of NCM in the left hemisphere yield a predictive value of future song learning success.

Most studies that aim at identifying the biological basis of song learning focus on the song control and auditory pathways. Intriguingly, none of the voxel-wise correlations identified in this study co-localize with any of the song control nuclei, even though evidence has been collected that, for example, specific microstructural properties of premotor cortical nucleus HVC are shaped by tutor song experience(20), and the functional properties of local inhibitory circuitries are tuned by the learning process(21), even after 60 dph. Both phenomena were not picked up by the voxel-wise correlations. Furthermore, the gross anatomy (volume) of the song control system nuclei reaches adult proportions by 60 dph(22). These findings as well as the current study suggest that the late sensorimotor and post critical period song refinement also involves areas that are not necessarily part of, but link to the song control system.

The structural architecture of left NCM measured during the sensory phase is able to predict future good or bad imitation outcome. This observation aligns with its previously established functional role in memorizing the tutor song during the sensory phase. More specifically, selective pharmacological blockage of the ERK or mTOR signaling cascades in the NCM of juvenile male zebra finches during exposure to the tutor prevents birds from memorizing and producing a tutor-like song in adulthood(23, 24). Tutor-song evoked immediate early gene (IEG) expression levels appear stronger in the left compared to the right NCM and song similarity correlates with the extent of lateralized IEG expression(25). Intriguingly, these data remind to left-ward lateralization of speech and language processing in humans(26), corroborate with a recently discovered asymmetry in tissue microstructure–measured by diffusion MRI– in the planum temporale related to (auditory) speech processing in human subjects(27), and connect tightly to our observation that mainly the left rather than the right NCM presents a correlation between tissue microstructure and tutor song imitation accuracy.

The difference between future good and bad learning outcome appears already written in the left NCM microstructure at 20 dph, before learning is known to commence(19). This finding might suggest that song learning starts earlier, that preparations of the neural substrate necessary for sensory learning occur before 20 dph, or that this structural signature is part of a natural predisposition to become a good or bad learner. Given that we observe a clear effect of tutor identity on final song learning accuracy –in line with others(4)– and that juvenile birds were randomly assigned to either their biological father or a foster father – therefore eliminating potential heritable effects of song learning success– the latter suggestion is less likely.

While the structural properties of left NCM already appear to differentiate between future good and bad performing birds in the sensory phase, the local volume and/or intrinsic tissue properties of the VP and tFA are not different between birds with a differential future learning outcome. Furthermore, the local volume of the CM and VP decreases when birds progressively become better at singing the tutor song. These findings suggest that the structural properties of the CM, VP and tFA are shaped during the sensorimotor phase.

Sensorimotor song refinement requires mechanisms that link motor commands with the associated sensory feedback such that it enables to detect and correct errors in own performance(28). Recent evidence appoints an important role to respectively the secondary auditory area CM and dopaminergic midbrain nuclei that synapse onto the vocal basal ganglia in these processes(29, 30). More specifically, the CM is reciprocally connected with NCM(31) and presents clear song-selective responses(32). A specific sub-field within the CM, the nucleus Avalanche (Av) exhibits singing-driven IEG expression that correlates with the amount of singing in normally hearing and in deafened birds(33). Av-projecting HVC neurons convey premotor signals to the Av and genetic ablation of HVC_Av_ neurons after sensory learning significantly impairs sensorimotor learning in juvenile male zebra finches(34). Furthermore, specific neuronal populations in the lateral portion of the CM, potentially co-localizing with the Av, selectively respond to white-noise played over song during singing but not when males are listening to their own song playback(30). Such signals might point to motor-to-auditory forward signals, like those observed in the human auditory cortex along speech production(35). In sum, its song selective neural responses(32), its reciprocal connectivity with NCM(31), and premotor input from HVC(34) that might participate in the generation of error-detection signals(30), set the CM as prime target capable of comparing own song performance towards pre-set performance goals such as the tutor song. Our findings complement these previous reports by showing that besides the functional properties, also the structural features of the CM are shaped throughout the sensorimotor learning phase.

Using optogenetic neuromodulation, two recent studies found mechanistic evidence that dopaminergic Area X projecting VTA neurons are indispensable for sensorimotor song learning in ontogeny(29). Furthermore, axonal collaterals of the DLM-projecting Area X cells –that make up the basal ganglia-thalamic component of the song control system– project to the VP where they synapse onto VTA/SNc projecting neurons(36). As such, the anterior forebrain pathway loops back onto its dopaminergic modulator. Besides carrying dopaminergic projections, the VP contains cholinergic neurons as well. Neurons located in the VP send cholinergic projections to two cortical regions responsible for the motor aspects of singing, i.e. HVC and RA(37) (Supplementary Fig. SI-1), and are capable of suppressing HVCs’ neural responses to birds’ own song by manipulation of the cholinergic projection neurons originating from the VP(38). Important to note is that the cholinergic projections in the VP are relatively sparse compared to the abundant dopaminergic projections and these cholinergic projection neurons co-localize with DLM-projecting Area X collaterals that terminate in the VP(15). This co-localization might point to a potential pathway via which descending auditory input affects HVC and RA. In sum, the VP appears an important integration center, where several pathways converge and perhaps form a closed loop system where dopaminergic midbrain nuclei can affect the basal ganglia to affect song output and vice versa, or cholinergic projections affect premotor cortical nuclei HVC and RA, based on error-signals originating from upstream auditory cortices(39). Our findings clearly complement functional studies by showing that the volume and most probably the organization of fibers of passage becomes rearranged when birds achieve better song copying accuracy. The exact role of each of these systems and the contribution of the VP to sensorimotor song imitation, however, requires further in-depth studies.

Several studies point to the importance of multi-sensory information for successful song learning and point to the nidopallium and arcopallium as potential integration centers(40). The cluster we observe in the tFA potentially extends on this multi-sensory hypothesis. The tFA carries projections from the basorostral nucleus in the rostral forebrain to the lateral arcopallium and caudolateral nidopallium. More specifically, the basorostral nucleus sends somatosensory and auditory information to the lateral arcopallium(14), which in turn projects to jaw premotor neurons and vocal and respiratory effectors(41). Also in humans, somatosensory information from facial skin and muscles of the vocal tract is vital for proper perception and production of speech(42, 43). Alternatively, these somatosensory and auditory descending projections may serve to control beak movements that modulate gape size during singing, as gape size can affect the acoustic properties of individual syllables(44). Taken together, even though this tract is not considered part of the traditional song control system, it might carry neuronal projections that are necessary for proper adjustment of vocalizations and be strengthened by vocal practicing.

In conclusion, the present findings clearly illustrate that song becomes better through practice and this appears associated with the structural properties of the CM, VP and tFA. However, how good the final song will be might depend heavily on template encoding in the secondary auditory area NCM at the early start of the sensory learning phase. Even though MRI cannot unambiguously identify the neurobiological mechanisms responsible for the structural signatures detected *in vivo*, nor can it establish causal relationships via correlation analyses, it provides a three-dimensional spatial map that can guide future studies to disentangle the exact contribution of each of these areas to song learning up to the cellular and molecular level. Such studies will be vital to understand the exact role of the VP and tFA in song learning. Instead of being purely a relay station, perhaps the VP exerts a more direct role as modulatory hub(45) as it receives, sends and relays dopaminergic and cholinergic projections to the vocal basal ganglia but also to the cortical components of the song control circuitry. Understanding the structural relationship between song learning accuracy and the tFA and the brain areas it connects might bring us a step closer in understanding how social interactions or sensory feedback help promote song learning success. These insights will have important consequences for identifying how neural networks are affected or can be modulated in disorders of speech. Furthermore, they clearly indicate that certain determinants for future success of performance appear to be set already at the earliest stages of the learning process, well before being actively engaged into practicing.

### Animals and Ethics statement

Male zebra finches (n = 16; *Taeniopyiga guttata*), bred in the local animal facility and were housed in individual cages together with an adult male (tutor), an adult female and one or two other juvenile zebra finches. At around 10 dph the juvenile birds were randomly assigned to an adult couple. This way some birds were co-housed with their biological parents, while others were raised by foster parents. Each cage was shielded from its neighboring cages so that the juvenile birds could hear all other birds of the room (6-12 other tutors and many other juveniles), but could interact (visual and auditory) with only one adult male bird. Juvenile birds will prefer to copy the song of the adult male bird with whom they can interact with(46). The ambient room temperature and humidity was controlled, the light-dark cycle was kept constant at 12h-12h, and food and water was available *ad libitum* at all times. In addition, from the initiation of the breeding program until the juvenile birds reached the age of 30 days post hatching (dph) egg food was provided as well. The Committee on Animal Care and Use at the University of Antwerp (Belgium) approved all experimental procedures (permit number 2012-43 and 2016-05) and all efforts were made to minimize animal suffering.

### Experimental design

#### DESIGN

We obtained MRI data of each bird during the sensory phase (20 and 30 dph), sensorimotor phase (40 and 65 dph), crystallization phase (90 and 120 dph) and one last time point well beyond the critical period for song learning (200 dph; Supplementary Fig. SI-1). Each imaging session, we collected a 3D anatomical scan and 2D Diffusion Tensor Imaging (DTI) data to evaluate respectively gross neuro-anatomy (volumetric analyses) and alterations to white matter tracts or intrinsic tissue properties. Starting from the advanced sensorimotor phase (i.e. 65 dph), we recorded the songs sung by the juvenile males the first day after each imaging session.

### Song recordings and analyses

To quantitatively evaluate the progression of sensorimotor learning and song refinement in male birds, we analyzed the first 20 (undirected) songs sung in the morning after ‘lights on’ in Sound Analysis Pro (SAP(3); http://soundanalysispro.com/). The undirected songs sung by the juvenile and adult male zebra finches and tutors were recorded in custom-build sound attenuation chambers equipped with the automated song detection setup implemented in SAP. All song analyses were performed off-line and calls and introductory notes were omitted from all analyses. First, the motif length (ms) of the first 20 songs sung during the morning (starting from the initiation of the photophase) was measured after which each individual motif was manually segmented into its different syllables based on sharp changes in amplitude and frequency. The latter measure was chosen to avert inconsistent determination of the syllable ending caused by more silent singing towards the last part of the syllable. Second, several acoustic features that reflect the spectro-temporal structure of individual syllables were quantified, i.e. (1) pitch-related measures that inform on the perceived tone of sounds (including pitch, mean frequency, peak frequency and goodness of pitch), (2) Wiener entropy that quantifies the tonality of sounds and is expressed on a logarithmic scale where white noise approaches ‘0’ and pure tones are characterized by large, negative Wiener entropy values, (3) syllable and inter-syllable interval duration. Furthermore, to evaluate syllable feature variability over the different ages, the standard deviation, as an estimate for vocal variability(47), was defined for each acoustic property. Next, similarity to tutor song was measured between song motifs using an automated procedure in SAP that quantifies the acoustic similarity between two songs based on pitch, FM, AM, goodness of pitch and Wiener entropy(3). Song similarity was calculated using the default settings of SAP (asymmetric comparisons of mean values, minimum duration 10ms, 10×10 comparisons), and % similarity was used for statistical testing. Further, according to the method conceptualized by Scharff and Nottebohm, motif sequence stereotypy was computed, based on visual assessment of sequence consistency and linearity(47). Sequence linearity reflects how consistent notes are ordered within the song motif by counting the different transition types of each syllable of the motif. Sequence consistency quantifies how often a particular syllable sequence occurs over different renditions of a specific motif. Song sequence stereotypy is defined as the average of sequence linearity and sequence consistency.

### MRI data acquisition

All MRI data were acquired on 7 T horizontal MR system (PharmaScan, 70/16 US, Bruker BioSpin GmbH, Germany) and a gradient insert (maximal strength: 400 mT/m; Bruker BioSpin, Germany), combined with a quadrature transmit volume coil, linear array receive coil designed for mice, following a previously described protocol(6). First, the zebra finches were anaesthetized with isoflurane (IsoFlo®, Abbott, Illinois, USA; induction: 2.0-2.5%; maintenance: 1.4-1.6%). While anaesthetized, the physiological condition of the birds was monitored closely by means of a pressure sensitive pad placed under the chest of the bird to detect the breathing rate, and a cloacal thermistor probe connected to a warm air feedback system to maintain the birds’ body temperature within narrow physiological ranges (40.0 ± 0.2) °C (MR-compatible Small Animal Monitoring and Gating system, SA Instruments, Inc.). Next, we collected DTI data using a four-shot spin echo (SE) echo planar imaging pulse sequence with the following parameters: TE 22 ms, TR 7000 ms, FOV (20×15) mm^2^, acquisition matrix (105×79), in-plane resolution (0.19×0.19) mm^2^, slice thickness 0.24 mm, b-value 670 s/mm², diffusion gradient duration (δ) 4 ms, diffusion gradient separation (Δ) 12 ms, 60 unique non-collinear diffusion gradient directions and 21 non-diffusion-weighted (b_0_) volumes. The entire DTI protocol was repeated twice to increase the SNR (total DTI scanning duration: 72 min). The field-of-view included the telencephalon and diencephalon which contain the auditory system and brain areas implicated in song control, the cerebellum and parts of the mesencephalon. Last, we collected a T_2_-weigthed 3D Rapid Acquisition with Relaxation Enhancement dataset with these settings: TE 55 ms, TR 2500 ms, RARE factor 8, FOV (18×16×10) mm^3^, matrix (256×92×64) zero-filled to (256×228×142), spatial resolution (0.07×0.17×0.16) mm^3^ zero-filled to (0.07 × 0.07 × 0.07) mm^3^, scan duration 29 min. The FOV of the 3D scan encapsulated the entire zebra finch brain. The entire scanning protocol took no longer than 2.5h. All animals recovered uneventfully after discontinuation of the anesthesia.

### MRI data processing

We processed both DTI and 3D RARE scans for voxel-based analyses using the following software packages: Amira (v5.4.0, FEI; https://www.fei.com/software/amira-3d-for-life-sciences/), ANTs (Advanced Normalization tools; http://stnava.github.io/ANTs/), FSL (FMRIB Software Library; https://fsl.fmrib.ox.ac.uk/fsl/fslwiki/FSL), and SPM12 (Statistical Parametric Mapping, version r 6225, Wellcome Trust Centre for Neuroimaging, London, UK, http://www.fil.ion.ucl.ac.uk/spm/) equipped with the Diffusion II toolbox (https://sourceforge.net/projects/diffusion.spmtools.p/) and DARTEL tools(48).

#### Deformation-Based Morphometry

First, we masked the individual 3D RARE scans (Amira 5.4.0) so that the datasets only include brain tissue. Second, we used the serial longitudinal registration (SLR) toolbox embedded in SPM12 to create one average ‘within-subject’ 3D dataset for each animal based on the individual masked 3D RARE scans acquired at the different ages(49). The SLR generated an average ‘within-subject’ 3D (‘midpoint average’) and jacobian determinant (j) maps. The latter maps are derived from the deformation field that contains the spatial transformation that characterizes the warp between the midpoint average and each individual time point image, and encode for each voxel the relative volume at a specific age with respect to the midpoint average. Next, the midpoint averages of all animals were inputted in the ‘buildtemplateparallel’ function of ANTs to create a between-subject population-based template. We used this initial ‘between-subject’ template to extract tissue probability maps reflecting mainly grey matter, white matter, or cerebrospinal fluid using the FMRIB Automated Segmentation Tool (FAST(50)) embedded in FSL. The three probability maps created in this step were used as tissue class *priors* for segmenting all individual midpoint averages using the default settings of the (old)segment batch in SPM12. The resulting tissue segments of the midpoint averages were used to create a segment-based template in Diffeomorphic Anatomical Registration through Exponentiated Lie Algebra (DARTEL)(48). Next, the ANTs-based T_2_-weighted between-subject template was warped via the ‘DARTEL: existing template’ batch to spatially match with the segment template and this template was used as anatomical reference space for all voxel-based analyses (referred to as ‘population-based template’). The jacobian determinant maps were warped to the reference space using the flow fields produced by DARTEL (with modulation to preserve relative volume differences existing between different subjects). Lastly, the warped modulated jacobian determinant maps were log-transformed and smoothed using a Gaussian kernel with FWHM (0.14×0.14×0.14) mm^2^.

#### Diffusion Tensor Imaging

The DTI data were pre-processed in the Diffusion II toolbox embedded in SPM12. First, we realigned the DTI volumes to correct for subject motion following a two-step procedure: an initial estimation based exclusively on the b_0_ images was followed by a linear registration including all (b_0_ and diffusion-weighted) volumes. Second, we co-registered the realigned DTI volumes to the individual 3D dataset acquired at the same age using normalized mutual information as objective function for inter-modal within-subject registration. In parallel, each individual masked 3D RARE dataset was bias corrected and spatially normalized to the population-based template using a 12-parameter affine global transformation followed by nonlinear deformations. These estimated spatial normalization parameters were applied to the co-registered DTI volumes so that the DTI data spatially matched the population-based template space. During this writing step, the diffusion data were upsampled to an isotropic resolution of 0.19 µm. In parallel, the diffusion vectors were adapted to account for potential (linear) rotations incurred by the realignment, co-registration and normalization procedures using the ‘copy and reorient diffusion information’ tool of the Diffusion II toolbox. Then, the diffusion tensor model was applied to the diffusion-weighted and b_0_-data to estimate the diffusion tensor and Eigenvalues (λ_1_, λ_2_, λ_3_). The Eigenvalues represent the principle axes of the radii of the 3D diffusion ellipsoid. Based on the Eigenvalues, the Fractional Anisotropy (FA) maps were computed.

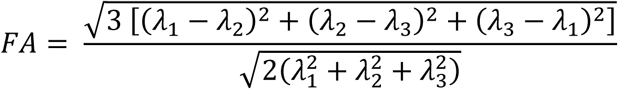

Several quality controls were performed throughout the dat acquisition and processing procedures. Those included a visual inspection for ghosting, excessive movement, and spatial correspondence of registered images to the reference space (after co-registration and spatial normalization procedures). Last, the DTI parameter maps were smoothed using a Gaussian kernel with FWHM of (0.38×0.38×0.38) mm^3^.

### Statistical analyses

Even though the MRI data were thoroughly checked at several stages in the pre-processing, an additional quality control was performed based on outlier detection. This analysis identified one outlier which, upon visual inspection of the datasets, appeared driven by abnormal cerebellar folding patterns (leading to suboptimal subsequent image registration) and was therefore excluded from all analyses. Furthermore, technical issues with the recordings of one tutor caused that we were not able to quantify song similarity to tutor song of two juvenile birds (one being the MRI-based outlier). Other technical issues lead to the loss of song recordings of two juvenile birds at 90 dph. This leaves 54 data points for voxel-based statistical testing (12 juveniles with 4 time points and 2 juveniles with 3 time points).

#### Song analyses

To analyze whether the song parameters change from 65 to 200 dph, we set up mixed-effect models including age as fixed effect, subject as random effect, subject*age as random slope and –only for the syllable level– syllable identity as random effect nested within subject. Furthermore, to determine whether song % similarity was dependent on the tutor by which the birds were raised, a mixed-effect model was performed with ‘tutor identity’ as fixed effect and ‘bird identity’ as random effect. We used the restricted maximum likelihood method to fit the data and assessed significance using F-tests with Kenward-Roger approximation. If a significant main effect could be observed for any of the song features examined, Tukey’s HSD (Honest Significant Difference) *post hoc* tests were performed to situate when in time actual changes occur.

#### Voxel-based statistical correlation analyses

The structural properties of the brain are tuned by learning and behavioral performance(8). To explore potential relationships between robust measures of song performance and the structural properties of the songbird brain, voxel-wise multiple regressions were set up between the smoothed MRI parameter maps (log mwj and FA maps) and % similarity or sequence stereotypy. When searching for correlations between local tissue volume and the song features, total brain volume was added to the statistical design as additional covariate. Unless explicitly stated, only clusters that survived a family-wise error (FWE) correction thresholded at *p*_*FWE*_<0.05 (or stricter) combined with a minimal cluster size (k_E_) of at least 5 or 20 voxels respectively for DTI and volume analyses, were considered significant. In contrast with song similarity, no supra-threshold voxels could be observed between any combination of smoothed MRI parameter maps and song sequence stereotypy. Clusters detected by the voxel-based correlation analyses were converted to ROIs (termed ‘cluster-based ROIs’, conversion at *p*_*uncorrected*_<0.001 k_E_≥40 voxels) of which the mean DTI metrics or modulated log-transformed jacobian determinant were extracted for *post hoc* statistical testing.

#### Cluster-based ROI analyses

The voxel-wise multiple regression in SPM does not allow including a random effect for bird identity. Consequently, by inserting repeated-measures data we violate the assumption of independency of measures. To correct for this potential confound, we performed two additional tests on the cluster-based ROI data. Firstly, we employed a repeated-measures correlation analysis(18) in Rstudio (version 1.1.383, Rstudio®, Boston, MA, USA; http://www.rstudio.com/). This latter test informs on the existence of consistent within-subject correlations between the two variables. Hence, this analysis informs on potential song learning-related structural changes in the brain. Secondly, Spearman’s ρ was calculated to test potential sources of between-bird variance in driving the correlations observed by the voxel-based analyses. Spearmans’ ρ was computed in JMP® (Version 13, SAS Institute Inc., Cary, NC, 1989-2007).

To explore the possibility of predicting future song learning outcome (good or bad), we ran a mixed-effect model analyses on the cluster-based ROI parameters extracted from MRI data obtained at 20, 30 and 40 dph. Bird identity was added a random factor, and age (20, 30, 40 dph) and learning outcome (good, bad) were included as fixed effects in the model. The restricted Maximum Likelihood method was used to fit the data and significance was assessed using *F*-tests with the Kenward-Roger approximation.

All additional tests on the cluster-based ROIs were corrected for multiple comparisons using false discovery rate (FDR) based on the Benjamini-Hochberg procedure(51). In brief, this method ranks all *p*-values from smallest (rank *i*=1) to largest (rank *i*=n). For each rank, the Benjamini-Hochberg critical value, (*i*/*m*)*Q*, is calculated. Where *i* is the rank, *m* is the total number of tests and *Q* is the false discovery rate set at 0.05. All *p*-values smaller than and including the largest *p*-value where *p*<(*i*/*m*)*Q* were considered significant.

The outcomes of all statistical tests performed in this study are summarized in the Supplementary Data 1-4.

## Acknowledgments

We thank Dr S. C. Woolley and Dr D. Vallentin for valuable discussions of the data and reading of earlier versions of the manuscript. The computational resources and services used in this work to build the population-based template were provided by the HPC core facility CalcUA of the Universiteit Antwerpen, the VSC (Flemish Supercomputer Center), funded by the Hercules Foundation and the Flemish Government – department EWI.

## Author contributions

JH conceptualized the study, acquired and analyzed the MRI data, and performed statistical analyses on the processed song data. KL analyzed the song data. MV advised on the statistical methodology and MRI data interpretation. AvDL conceptualized the study, obtained funding resources and supervised the whole project. All the authors discussed the results, contributed to writing the manuscript, and approved the final version of the manuscript for submission.

## Data and materials availability

The data related to this paper can be requested to the corresponding author.

## Supplementary Materials

### Supplementary Figure SI-1: Schematic overview of the songbird brain and pathways important for vocal learning, and study design

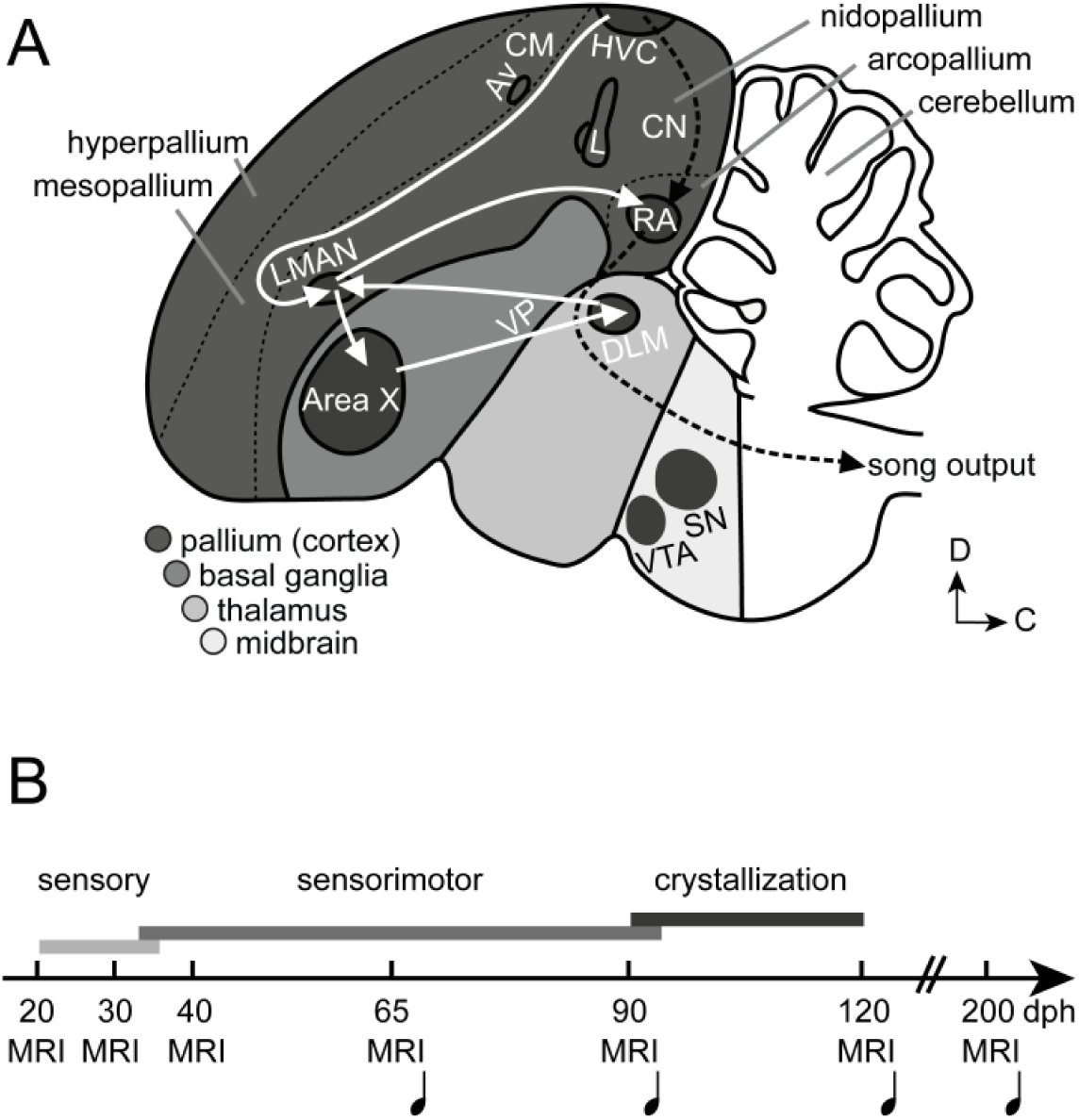
Schematic overview of the male zebra finch brain (A) and study design (B). **A)** The arrows connect different components of the song control circuitry, i.e. a set of interconnected nuclei responsible for singing that consists of two major pathways. The white arrows illustrate the anterior forebrain pathway (AFP) including vocal basal ganglia important for introduction variability in vocal output during sensorimotor vocal practicing, the black dashed arrows represent the caudal motor pathway (CMP) responsible for the motor aspect of singing. The CMP consists of a direct connection between premotor nucleus HVC (abbreviation used as a proper name;) and the robust nucleus of the arcopallium (RA). HVC premotor neurons synapse onto RA neurons which in turn sends myotopically organized projections onto motor nuclei in the brainstem (of the vocal organ) and to respiratory nuclei which are capable of modifying the acoustic features of vocal sounds. The AFP forms an indirect connection between HVC and RA by sequentially connecting HVC, Area X, the dorsolateral nucleus of the medial part of the anterior thalamus (DLM), the lateral magnocellular nucleus of the anterior nidopallium (LMAN), a frontal cortical nucleus and RA. Area X is a large brain region that is functionally and structurally similar to the mammalian basal ganglia. The ventral tegmental area (VTA) and substantia nigra (SN) are dopaminergic midbrain nuclei that send dopaminergic projections to Area X. Field L, the caudomedial nidopallium (medial part of the CN) and caudal mesopallium (CM), which includes the nucleus Avalanche (Av), are part of the auditory system. **B)** Structural MRI data were acquired during the sensory phase (20, 30 dph), sensorimotor phase (40, 65 dph), crystallization phase (90, 120 dph) and well after song is mastered (200 dph). During each imaging session, we collected a DTI and 3D anatomical scan. The first days after each imaging session, we recorded the song sung by the juvenile male birds (♪), starting from 65 dph. Abbreviations: D: dorsal; VP: ventral pallidum.

### Supplementary Data 1: Song analyses

The duration of the inter-syllable intervals gradually shortens from sensorimotor to song crystallization and even after song crystallization towards 200 dph (*p*<0.0001 *F*_*(3,38.2)*_=13.8789; Figure SI-2 A). This indicates that birds gradually sing faster. Syllable Wiener entropy scores decrease during the sensorimotor phase (*p*=0.0032 *F*_*(3,37.8)*_=5.48; Figure SI-2 B), meaning that the syllables gain in tonality. None of the pitch-related measures, or frequency and amplitude modulation changed over time (Table SI-1-A). Further, by quantifying the standard deviation of the spectro-temporal features, we observe that syllables are sung with lower acoustic variability when song crystallizes between 90 and 120 dph (Figure SI-2 C).

**Table SI-1-A:** Summary of the mixed-effect model analyses on the song scores.

| Syllable features (Main effect of age) |  |
| --- | --- |
| inter-syllable interval duration | $p<0.0001$ $F_{(3,38,2)}=13.8789$ |
| syllable duration | $p=0.5534$ $F_{(3,37,7)}=0.7078$ |
| Wiener entropy | $p=0.0032$ $F_{(3,37,8)}=5.4803$ |
| pitch | $p=0.8897$ $F_{(3,37,7)}=0.2088$ |
| mean frequency | $p=0.1289$ $F_{(3,38,6)}=2.0078$ |
| peak frequency | $p=0.0649$ $F_{(3,38,6)}=2.6144$ |
| goodness of pitch | $p=0.2211$ $F_{(3,38,1)}=1.5348$ |
| FM | $p=0.3279$ $F_{(3,37,4)}=1.1867$ |
| AM | $p=0.5470$ $F_{(3,37,4)}=0.7189$ |
| Standard deviation of syllable features (Main effect of age) |  |
| inter-syllable interval duration | $p<0.0001$ $F_{(3,25,4)}=20.8043$ |
| syllable duration | $p=0.0064$ $F_{(3,34,3)}=4.8560$ |
| Wiener entropy | $p=0.0021$ $F_{(3,36,9)}=5.9192$ |
| pitch | $p=0.5824$ $F_{(3,41,4)}=0.6583$ |
| mean frequency | $p=0.0132$ $F_{(3,37,5)}=4.0891$ |
| peak frequency | $p=0.0280$ $F_{(3,38,1)}=3.3787$ |
| goodness of pitch | $p=0.0621$ $F_{(3,33,1)}=2.6910$ |
| FM | $p=0.0700$ $F_{(3,33,2)}=2.5803$ |
| AM | $p=0.0982$ $F_{(3,41,5)}=2.2367$ |
| Motif features (Main effect age) |  |
| song sequence stereotypy | $p=0.0052$ $F_{(3,38,4)}=4.7904$ |
| % similarity | $p=0.0251$ $F_{(3,37,0)}=3.4890$ |
| Motif features (Main effect tutor) |  |
| song sequence stereotypy | $p=0.1389$ $F_{(7,7,3)}=2.3350$ |
| % similarity | $p=0.0159$ $F_{(7,6,1)}=6.7597$ |

To test for a main effect of age, a mixed-effect model ( $n = 14-16$ ; details in Methods section) was set up for each song score separately with age as fixed effect, subject as random effect, subject\*age as random slope and –only for statistical analyses on the syllable feature level– syllable identity as random effect nested within subject. To test for a main effect of tutor, a mixed-effect model was executed for song similarity or sequence stereotypy separately with tutor as fixed effect and subject as random effect. The restricted
Maximum Likelihood method was used to fit the data and significance was assessed using $F$ -tests with the Kenward-Roger approximation.

**Figure SI-2:**
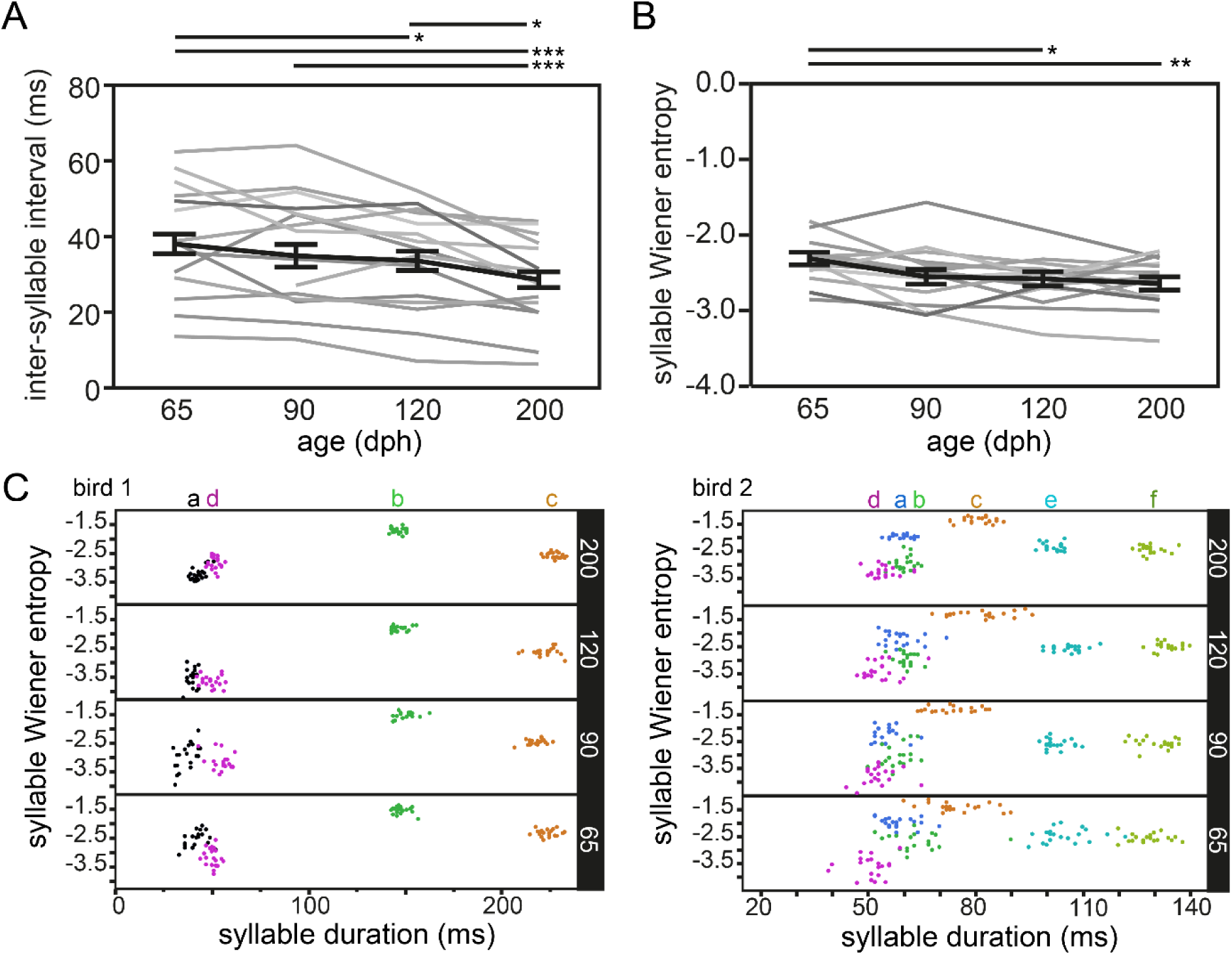
Song syllable scores. Each thin grey line (all graphs except for C) refers to the average performance of an individual bird over the different ages. The bold line presents the average group performance (mean ± s.e.m.; n=14; 20 data points per time point per bird). Graph **A** and **B** contain the syllable feature scores, resp. inter-syllable duration and syllable entropy. Both decrease significantly over the course of the study (mixed-effect model main effect of age: inter-syllable interval duration: *p*<0.0001 *F*_*(3,38.2)*_=13.8789; Syllable Wiener entropy: *p*=0.0032 *F*_*(3,37.8)*_=5.4803). **C** aims to visualize the change in standard deviation of specific syllables (each color presents the data of one syllable (indicated by letters) of two representative birds. Each horizontal part of the graph presents data obtained at one time point (top to bottom: 200, 120, 90 and 65 dph). The size of the cluster of syllables a and d in bird 1 and syllables a-b-c-d-e of bird 2 decreases markedly between 65 and 200 dph, indicative of smaller variation between different syllable renditions towards older ages. Asterisks indicate significant differences over time indicated by a mixed model analysis with *post hoc* Tukey’s HSD test. *: p<0.05; ** p<0.01; *** p<0.001. The previously detected main effect of tutoring bird on song similarity outcome did not seem to depend on the length of the songs, as based on similarity scores obtained at 200 dph no differences in the number of syllables for birds having a ‘high’ (>68*) or ‘low’ (<68*) similarity score could be observed (Table SI-1-B).

**Table SI-1-B:** Number of syllables in the songs of good and bad learners.

| > 65-68% similarity |  | < 65-68% similarity |  |
| --- | --- | --- | --- |
| Bird ID | Motif structure | Bird ID | Motif structure |
| 03 | ABCD | 05 | ABCD |
| 17 | ABCDEF | 07 | ABCDE |
| 18 | ABCDEF | 09 | ABCDE |
| 20 | ABCD | 10 | ABCDE |
| 21 | ABC | 35 | ABC |
| 23 | ABCD |  |  |
| 31 | ABC |  |  |

‘> 65-68 %’ indicates that the birds always sung song copies with a song similarity to tutor song score of at least 65-68%, while ‘< 65-68%’ indicates that birds always sung songs with song similarity score lower than 65-68% similarity to tutor song. The 65-68% threshold was chosen arbitrarily based on the overall performance of the birds within the study.

### Supplementary Data 2: Voxel-based multiple regression

**Table SI-2-A:** Summary of the voxel-based multiple regressions (% similarity and FA or logmwj).

| Correlation between | Cluster | Hemisphere | Cluster |  | Peak |  |
| --- | --- | --- | --- | --- | --- | --- |
| | | | $k_E$ | $p_{FWE}$ | $T$ | $p_{FWE}$ |
| % similarity and FA | tFA | Left | 13 | 0.002 | 6.81 | <0.001 |
|  |  | Right | / | 0.034 | 6.42 | 0.001 |
|  | NCM | Left | 5 | 0.001 | 5.69 | 0.019 |
|  |  | Right | / | 0.024 | 5.55 | 0.032 |
|  | VP |  | 479 | <0.001 | 6.38 | 0.002 |
| % similarity and log mwj | VP |  | 2022 | <0.001 | 8.06 | <0.001 |
|  | CM | Left | 1701 | <0.001 | 7.10 | 0.001 |
|  |  | Right | 3465 | <0.001 | 7.42 | <0.001 |

‘log mwj’ refers to the log-transformed, modulated and warped jacobian determinants; FA stands for
Fractional Anisotropy, one of the DTI metrics. This table summarizes the outcome of the voxel-based
multiple regression based on 54 data points (12 birds with 4 time points and 2 birds with 3 time points). The
‘Cluster’ and ‘Peak’ columns refer to two different levels of assessing significance, respectively cluster-
based inference and peak- or single voxel-based inference where the $T$ - and $p$ -value of the voxel with highest
significance of the cluster is reported. Only clusters surviving $p_{FWE}<0.05$ and $k_E>5$ voxels were considered
significant. Grey font refers to clusters that were only visible when exploring the data at an exploratory
statistical threshold, i.e. $p_{uncorrected}<0.001$ and $k_E>40$ voxels. At this lower statistical threshold for visual
assessment of the statistical maps, only clusters that still appeared significant with FWE correction at the

### Supplementary Data 3: Within-and between-bird correlation analyses of cluster-based ROIs

**Table SI-3-A:** Summary of within-and between-subject correlations of the cluster-based ROIs.

| Correlation between | Cluster-based ROI | Hemisphere | Rmcorr |  | 65 dph |  | 200 dph |  |
| --- | --- | --- | --- | --- | --- | --- | --- | --- |
| | | | r | p | Spearman's $\rho$ | p | Spearman's $\rho$ | p |
| % similarity and FA | tFA | Left | 0.1290 | 0.4200 | 0.7846 | 0.0009 | 0.7714 | 0.0012 |
|  |  | Right | 0.0215 | 0.8940 | 0.7099 | 0.0045 | 0.7143 | 0.0041 |
|  | NCM | Left | 0.2560 | 0.1060 | 0.6747 | 0.0081 | 0.6835 | 0.0070 |
|  |  | Right | 0.3880 | 0.0121 | 0.6396 | 0.0138 | 0.5692 | 0.0336 |
|  | VP |  | 0.4960 | 0.0010 | 0.8154 | 0.0004 | 0.7890 | 0.0008 |
| % similarity and log mwj | VP |  | -0.4290 | 0.0057 | -0.7978 | 0.0006 | -0.4418 | 0.1138 |
|  | CM | Left | -0.3910 | 0.0126 | -0.5297 | 0.0514 | -0.3055 | 0.2882 |
|  |  | Right | -0.4160 | 0.0075 | -0.4769 | 0.0846 | -0.1912 | 0.5126 |

'log mwj' refers to the log-transformed, modulated and warped jacobian determinants; FA stands for Fractional Anisotropy, one of the DTI metrics. 'r' is the repeated-measures correlation coefficient of the within-subject correlation analyses. Spearman's $\rho$ informs on potential correlations between the MRI parameters and song similarity at a specific time point between birds. Abbreviations: dph: days post hatching.

**Table SI-3-B:** Benjamini-Hochberg FDR correction for multiple comparisons of rmcorr analyses.

| MRI parameter | Cluster-based ROI | Hemisphere | p value | rank | $(i/m)Q$ |
| --- | --- | --- | --- | --- | --- |
| FA | VP |  | **0.0010** | **1** | **0.0063** |
| Log mwj | VP |  | **0.0057** | **2** | **0.0125** |
| Log mwj | CM | Right | **0.0075** | **3** | **0.0188** |
| FA | NCM | Right | **0.0121** | **4** | **0.0250** |
| Log mwj | CM | Left | **0.0126** | **5** | **0.0313** |
| FA | tFA | Left | 0.4200 | 6 | 0.0375 |
| FA | tFA | Right | 0.8940 | 7 | 0.0438 |
| FA | NCM | Left | 0.1060 | 8 | 0.0500 |

FDR rate = 0.05; number of tests = 8; $i$ is the rank, $m$ is the total number of tests and $Q$ is the false
discovery rate set at 0.05. Only those tests that survive FDR correction for multiple comparisons are
highlighted **bold**. 'rmcorr' is the repeated-measures correlation analysis.

**Table SI-3-C:** Benjamini-Hochberg FDR correction for multiple comparisons of Spearmans’ ρ analyses.

| MRI parameter | Cluster-based ROI | Age | Hemisphere | *p* value | rank | (i/m)Q |
| --- | --- | --- | --- | --- | --- | --- |
| FA | VP | 65 |  | **0.0004** | **1** | **0.0031** |
| Log mwj | VP | 65 |  | **0.0006** | **2** | **0.0063** |
| FA | VP | 200 |  | **0.0008** | **3** | **0.0094** |
| FA | tFA | 65 | Left | **0.0009** | **4** | **0.0125** |
| FA | tFA | 200 | Left | **0.0012** | **5** | **0.0156** |
| FA | tFA | 200 | Right | **0.0041** | **6** | **0.0188** |
| FA | tFA | 65 | Right | **0.0045** | **7** | **0.0219** |
| FA | NCM | 200 | Left | **0.0070** | **8** | **0.0250** |
| FA | NCM | 65 | Left | **0.0081** | **9** | **0.0281** |
| FA | NCM | 65 | Right | **0.0138** | **10** | **0.0313** |
| FA | NCM | 200 | Right | **0.0336** | **11** | **0.0344** |
| Log mwj | CM | 65 | Left | 0.0514 | 12 | 0.0375 |
| Log mwj | CM | 65 | Right | 0.0846 | 13 | 0.0406 |
| Log mwj | VP | 200 |  | 0.1138 | 14 | 0.0438 |
| Log mwj | CM | 200 | Left | 0.2882 | 15 | 0.0469 |
| Log mwj | CM | 200 | Right | 0.5126 | 16 | 0.0500 |

FDR rate = 0.05; number of tests = 16; *i* is the rank, *m* is the total number of tests and *Q* is the false discovery rate set at 0.05. Only those tests that survive FDR correction for multiple comparisons are highlighted **bold**.

### Supplementary Data 4: Predict success future song learning outcome

**Table SI-4-A:** Summary of mixed-effect model.

| MRI parameter | Cluster-based ROI | Hemisphere | Interaction good-bad*age | Main effect good-bad |
| --- | --- | --- | --- | --- |
| FA | NCM | Left | $p=0.1146 F_{(2,19,6)}=2.4242$ | $p=0.0003 F_{(1,10,2)}=28.3681$ |
| | | Right | $p=0.0289 F_{(2,19,4)}=4.2797$ | $p=0.2550 F_{(1,10,1)}=1.4557$ |
| | tFA | Left | $p=0.1378 F_{(2,19,4)}=2.1989$ | $p=0.0998 F_{(1,10,1)}=3.2809$ |
| | | Right | $p=0.0113 F_{(2,19,2)}=5.7186$ | $p=0.1198 F_{(1,10,1)}=2.8898$ |
| | VP | | $p=0.5908 F_{(2,19,3)}=0.5409$ | $p=0.5856 F_{(1,10,0)}=0.3173$ |
| log mwj | VP | | $p=0.6713 F_{(2,20,0)}=0.4066$ | $p=0.0463 F_{(1,10,0)}=5.1686$ |
| | CM | Left | $p=0.1188 F_{(2,20,0)}=2.3740$ | $p=0.3280 F_{(1,10,0)}=1.0577$ |
| | | Right | $p=0.4105 F_{(2,20,0)}=0.9313$ | $p=0.5240 F_{(1,10,0)}=0.4359$ |

This table summarizes the results of the mixed-effect model analyses testing for an interaction between age (20-30-40 dph) and future good-bad learning outcome (n=14 birds). The mixed-effect model includes a fixed effect for age and for learning outcome (good-bad), and a random factor for bird identity. The restricted Maximum Likelihood method was used to fit the data and significance was assessed using *F*-tests with the Kenward-Roger approximation.

**Table SI-4-B:** Benjamini-Hochberg FDR correction for multiple comparisons of the interaction good-bad*age.

| MRI parameter | Cluster-based ROI | Hemisphere | *p* value | rank | (i/m)Q |
| --- | --- | --- | --- | --- | --- |
| FA | tFA | Right | 0.0113 | 1 | 0.0063 |
| FA | NCM | Right | 0.0289 | 2 | 0.0125 |
| FA | NCM | Left | 0.1146 | 3 | 0.0188 |
| Log mwj | CM | Left | 0.1188 | 4 | 0.0250 |
| FA | tFA | Left | 0.1378 | 5 | 0.0313 |
| Log mwj | CM | Right | 0.4105 | 6 | 0.0375 |
| FA | VP |  | 0.5908 | 7 | 0.0438 |
| Log mwj | VP |  | 0.6713 | 8 | 0.0500 |

FDR rate = 0.05; number of tests = 8; *i* is the rank, *m* is the total number of tests and *Q* is the false discovery rate set at 0.05; None of the tests survives FDR correction.

**Table SI-4-C:** Benjamini-Hochberg FDR correction for multiple comparisons of the main effect good-bad.

| MRI parameter | Cluster-based ROI | Hemisphere | *p* value | rank | (*i*/*m*)*Q* |
| --- | --- | --- | --- | --- | --- |
| FA | NCM | Left | **0.0003** | **1** | **0.0063** |
| Log mwj | VP |  | 0.0463 | 2 | 0.0125 |
| FA | tFA | Right | 0.0998 | 3 | 0.0188 |
| FA | tFA | Right | 0.1198 | 4 | 0.0250 |
| FA | NCM | Right | 0.2550 | 5 | 0.0313 |
| Log mwj | CM | Left | 0.3280 | 6 | 0.0375 |
| Log mwj | CM | Right | 0.5240 | 7 | 0.0438 |
| FA | VP |  | 0.5856 | 8 | 0.0500 |

FDR rate = 0.05; number of tests = 8; *i* is the rank, *m* is the total number of tests and *Q* is the false discovery rate set at 0.05. Only those tests that survive FDR correction for multiple comparisons are highlighted **bold**.

**Figure SI-3:**
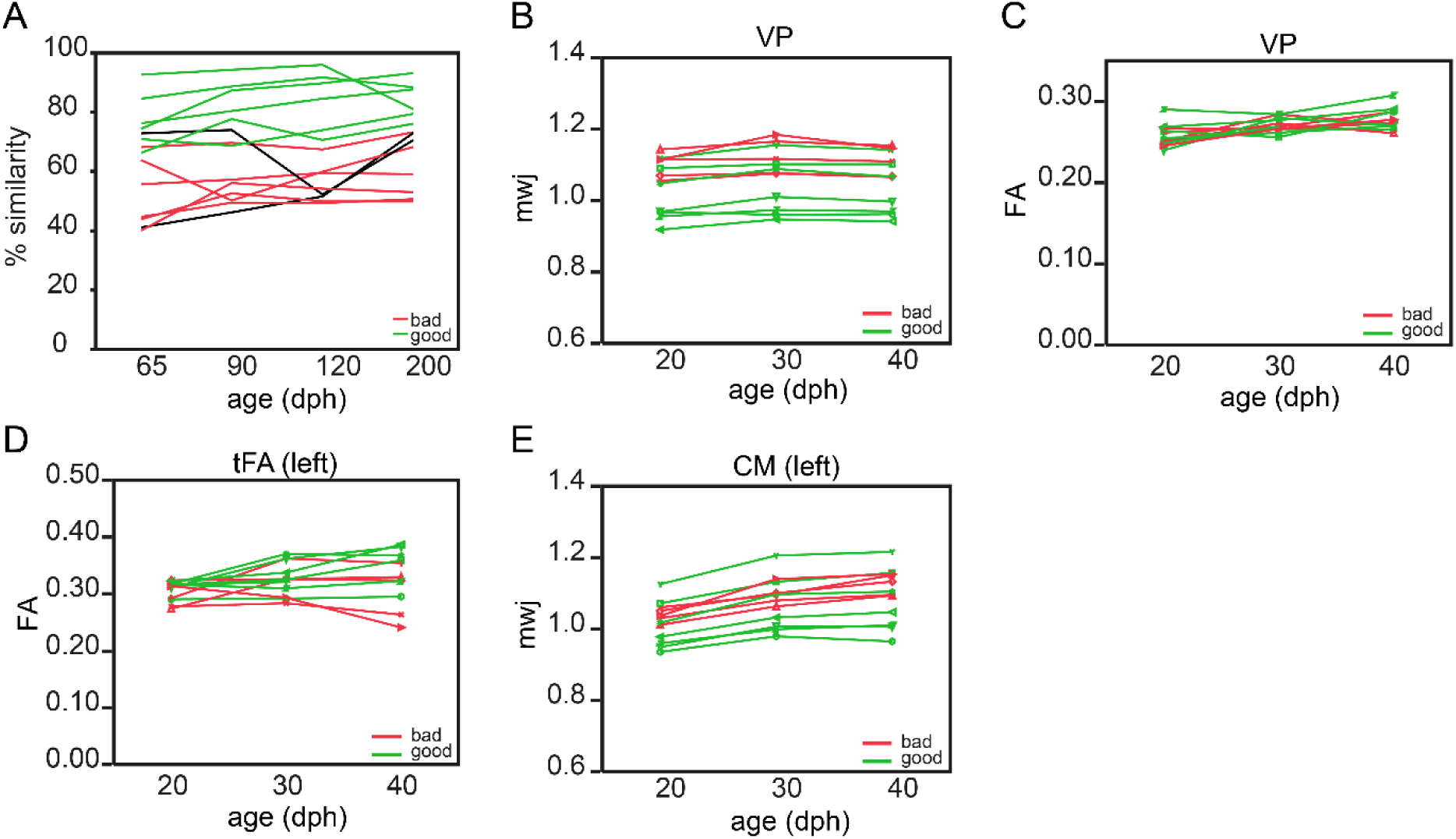
The structural properties of the VP, CM and tFA cannot predict future song learning accuracy. Graph A presents the learning curve of the good (green) and bad (red) learning birds from 65 to 200 dph. The two birds that could clearly traverse the 65-68% song similarity score could not be assigned to either group and are presented by a thin black line. (Details on the distinction between good and bad learners can be found in the Results section.) Graphs B-E present the difference in Fractional Anisotropy (FA) or mwj (local tissue volume) during the sensory (20-30 dph) or early sensorimotor phase (40 dph) of the cluster detected by the voxel-wise multiple regression between good (green) and bad (red) vocal learners. The outcome of the statistical tests is summarized in Tables SI-4-A, B, C.

